# NHX-type Na^+^(K^+^)/H^+^ antiporter activity is required for endomembrane trafficking and ion homeostasis in *Arabidopsis thaliana*

**DOI:** 10.1101/382663

**Authors:** Jonathan Michael Dragwidge, Stefan Scholl, Karin Schumacher, Anthony Richard Gendall

## Abstract

The regulation of ion and pH homeostasis of endomembrane organelles is critical for functional protein trafficking, sorting and modification in eukaryotic cells. pH homeostasis is maintained through the activity of vacuolar H^+^-ATPases (V-ATPases) pumping protons (H^+^) into the endomembrane lumen, and counter-action by cation/proton exchangers such as the NHX family of Na^+^(K^+^)/H^+^ exchangers. In plants, disturbing V-ATPase activity at the *trans*-Golgi network/early endosome (TGN/EE) impairs secretory and endocytic trafficking. However, it is unclear if the endosomal NHX-type antiporters NHX5 and NHX6 play functionally similar roles in endomembrane trafficking through maintaining ion and pH homeostasis. Here we show through genetic, pharmacological, and live-cell imaging approaches that double knockout of endosomal isoforms *NHX5* and *NHX6* results in impairment of endosome motility, protein recycling at the TGN/EE, but not in the secretion of integral membrane proteins. Furthermore, we report that *nhx5 nhx6* mutants are partially insensitive to osmotic swelling of TGN/EE induced by the monovalent cation ionophore monensin. Similarly, *nhx5 nhx6* cells are unresponsive to late endosomal swelling by the phosphatidylinositol 3/4-kinase inhibitor wortmannin, demonstrating that NHX5 and NHX6 are required for maintaining endosomal cation balance. Lastly, we report that the distal region of the cytosolic tail of NHX6 is required for mediating NHX6 localisation to late endosomes, but does not appear to be essential for NHX6 function.

## Introduction

The endomembrane system of eukaryotic cells is composed of a complex series of interconnected compartments which function in the synthesis, sorting, transport and degradation of proteins. For these cellular processes to operate efficiently, endomembrane organelles must control their luminal pH by balancing the activity of proton pumps and cation channels (Casey et al., 2010). In plants the *trans*-Golgi network also acts as an early endosome, and functions to sort and transport both newly endocytosed and secretory proteins (Viotti et al., 2010). The TGN/EE maintains a large proton gradient for functional protein sorting and secretion which is achieved through V-ATPase mediated proton pumping into the TGN/EE lumen (Schumacher, 2014). This acidification results in the TGN/EE as the most acidic plant endomembrane compartment (Martinière et al., 2013; Shen et al., 2013). Conversely, cation/proton exchangers including the NHX family of Na^+^,K^+^/H^+^ exchangers and CHX family of K^+^/H^+^ exchangers act as a proton leak to alkalinise the lumen of endosomes and assist in fine tuning organelle pH (Bassil et al., 2012; Brett et al., 2005b; Chanroj et al., 2012). Moreover, cation/proton exchangers also function in cation detoxification and are important for salt tolerance in plants (Rodríguez-Rosales et al., 2009).

Intracellular NHX-type exchangers have evolutionarily conserved roles in ion and pH homeostasis, and in plants also function in development and in protein trafficking to the vacuole (Bassil et al., 2012; Chanroj et al., 2012; Dragwidge et al., 2018). *Arabidopsis thaliana* has eight NHX genes; four encode tonoplast localised proteins (AtNHX1-AtNHX4), two are endosomal-localised (AtNHX5-AtNHX6), and two are present on the plasma membrane (AtNHX7/SOS1 and AtNHX8) (Brett et al., 2005b). In *A. thaliana* double knockouts of the endosomal NHX isoforms *nhx5 nhx6* show defects in vacuolar transport, with delayed trafficking of the endocytic tracer dye FM4-64 and mis-secretion of a vacuolar targeted yeast carboxypeptidase-Y (CPY) fragment (Bassil et al., 2011a). Furthermore, in embryos *nhx5 nhx6* mutants have defects in processing and transport of seed storage proteins to the vacuole (Ashnest et al., 2015; Reguera et al., 2015). Similar vacuolar trafficking defects have been described in yeast, with the knockout of the single endosomal *nhx1* gene causing altered CPY secretion, delayed vacuolar trafficking, and defects in late endosome/ multi-vesicular body (LE/MVB) formation and sorting (Bowers et al., 2000; Brett et al., 2005a; Kallay et al., 2011; Mitsui et al., 2011). Furthermore, RNAi silencing of mammalian endosomal orthologs *NHE6* and *NHE8* leads to disruptions in endosome trafficking and recycling (Lawrence et al., 2010; Ohgaki et al., 2010), demonstrating that eukaryotic endosomal NHXs have a conserved role in subcellular protein trafficking and recycling.

In plants endosomal NHX antiporters have been implicated in the trafficking of soluble cargo proteins to the vacuole. Soluble proteins such as seed storage proteins are synthesised in the ER, bind vacuolar sorting receptors (VSRs) and transit towards the TGN/EE (Künzl et al., 2016), before budding and maturation of the TGN/EE into the LE/MVB which then ultimately fuse with the vacuole (Scheuring et al., 2011). In *A. thaliana nhx5 nhx6* mutants have inhibited VSR-cargo interactions and mis-processing of seed storage proteins (Ashnest et al., 2015; Reguera et al., 2015). These defects are believed to be caused by hyper-acidification of endomembrane luminal compartments in *nhx5 nhx6* knockouts. Conversely, reduction of V-ATPase activity at the TGN/EE through Concanamycin-A treatment or in the *det3* mutant results in alkalinisation of the TGN/EE, and leads to defects in recycling and secretion pathways, including delayed trafficking to the vacuole and reduced Golgi and TGN/EE motility (Dettmer et al., 2006; Luo et al., 2015; Viotti et al., 2010). These findings demonstrate that maintaining endomembrane luminal pH of the TGN/EE is essential for functional protein secretion, recycling, and vacuole transport. While the function of NHX5 and NHX6 in vacuolar trafficking of soluble cargo proteins has been well described, it has not been identified whether NHX5 and NHX6 activity is also necessary for functional secretion, sorting and recycling of integral membrane proteins.

Here we investigated the effects of disrupted endomembrane pH and ion balance through dissection of the secretory and endocytic transport pathways in the double knockout *nhx5 nhx6*. Through live cell-imaging we reveal that *nhx5 nhx6* mutants have reduced Golgi and TGN/EE motility and defects in the recycling of transmembrane receptors from the TGN/EE. Furthermore, our results reveal that *nhx5 nhx6* endosomes are insensitive to osmotic swelling induced by the ionophore monensin, or with the late endosome inhibitor wortmannin. Moreover, we identified that the distal region of the cytosolic tail of NHX6 is required for mediating NHX6 localisation to the late endosome.

## Results

### NHX5 and NHX6 are involved in endomembrane compartment motility

In plant cells the movement of endomembrane organelles through the secretory and endocytic pathways is essential for functional protein delivery, and is highly dependent on the cytoskeleton network of actin filaments and microtubules (Brandizzi and Wasteneys, 2013). As alkalinisation of the TGN/EE has been shown to reduce Golgi and TGN/EE motility (Luo et al., 2015), we questioned whether the hyper-acidification of endomembrane organelles in *nhx5 nhx6* could also negatively affect their motility. We generated stable Arabidopsis *nhx5 nhx6* lines expressing the endosomal markers YFP-Got1p (Golgi), and YFP-VTI12 (TGN/EE) and examined endosomal motility through live cell spinning disk confocal microscopy. Quantitative analysis revealed that the movement of Golgi and TGN/EE vesicles were significantly reduced in *nhx5 nhx6* cells compared to wild type (Fig. 1A-E, S1A-B). Additionally, the proportion of slower moving bodies (< 10 μm min^−1^) was more than two-fold higher in *nhx5 nhx6* compared to wild type, demonstrating that a high proportion of vesicles exhibited minimal movement in *nhx5 nhx6* (Fig. S1C). We also assessed the straightness of particle tracks to indirectly assess whether organelle behaviour, or their potential association with the cytoskeleton may be altered. Quantification revealed a significant reduction in the straightness of TGN/EE trajectories in *nhx5 nhx6* cells, indicating that these vesicles displayed more disordered, non-continuous movement (Fig. 1E), typical of cytoskeletal independent endosome movement (Akkerman et al., 2011).

**Figure 1.**
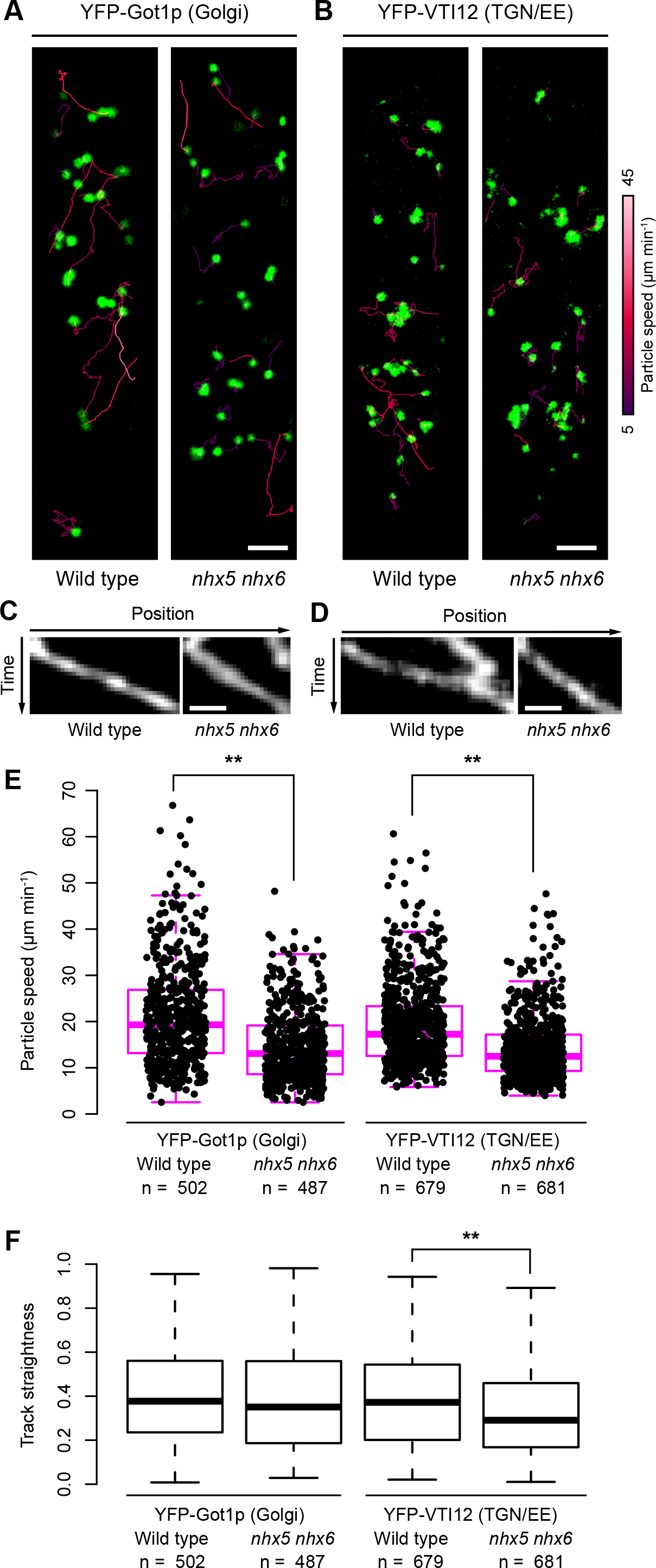
Golgi and TGN/EE motility are reduced in *nhx5 nhx6*. (A-B) Time-lapse spinning disk confocal microscopy of Golgi localised YFP-Got1p (A), and TGN/EE localised YFP-VTI12 (B) in hypocotyl cells from 4 day old dark grown Arabidopsis seedlings. The trajectory of particles visible from the initial frame of the time lapse are shown, coloured by mean particle speed. (C and D) Kymograph of a representative continuously moving Golgi (C) and TGN/EE (D) particle over 20 seconds. (E) Box plot of mean Golgi and TGN/EE particle speeds. ** p < 0.01, from Kolmogorov-Smirnov and Mann-Whitney tests. n = number of particle tracks analysed from ≥ 6 cells. (F) Box plot of Golgi and TGN/EE particle straightness, where 1 indicates a perfectly straight line. ** p < 0.01 from Kolmogorov-Smirnov and Mann–Whitney U tests. Scale bars = 5 μm (A and B), 2 μm (C and D). See also Movie S1 and Figure S1.

### BRI1 recycling, but not secretion is reduced in *nhx5 nhx6*

Next, we investigated whether the transport or recycling of transmembrane receptors at the TGN/EE may be inhibited in *nhx5 nhx6* cells. We employed the well characterised receptor kinase BRASSINOSTEROID-INSENSITVE 1 (BRI1) as it is constitutively endocytosed from the plasma membrane to the TGN/EE, where it is sorted for recycling back to the plasma membrane or degradation towards the vacuole (Dettmer et al., 2006; Geldner et al., 2007; Irani et al., 2012). We treated root cells expressing BRI1-GFP with the fungal toxin Brefeldin-A (BFA) to reversibly inhibit TGN/EE and Golgi trafficking (Geldner et al., 2001; Richter et al., 2007), and assessed the recycling of BRI1-GFP out of BFA bodies after washout (Fig. 2A). Quantification revealed that *nhx5 nhx6* cells had larger BFA bodies and a higher proportion of cells containing BFA bodies after washout (Fig. 2B - C). These results indicate that BRI1 recycling from the TGN/EE is impaired in *nhx5 nhx6*, and together with similar BRI1 recycling defects in *det3* mutants (Luo et al., 2015), suggests pH sensitive trafficking machinery are required for efficient BRI1 recycling. We also assessed growth response of *nhx5 nhx6* treated with the V-ATPase inhibitor Concanamycin A (ConcA) which disrupts TGN/EE structure and endocytic and secretory trafficking (Dettmer et al., 2006; Viotti et al., 2010). In *nhx5 nhx6* hypocotyl elongation was notably reduced compared to wild type and showed similar hypersensitivity to ConcA as *det3* (Fig. S2), demonstrating that *nhx5 nhx6* knockouts have impaired TGN/EE function.

**Figure 2.**
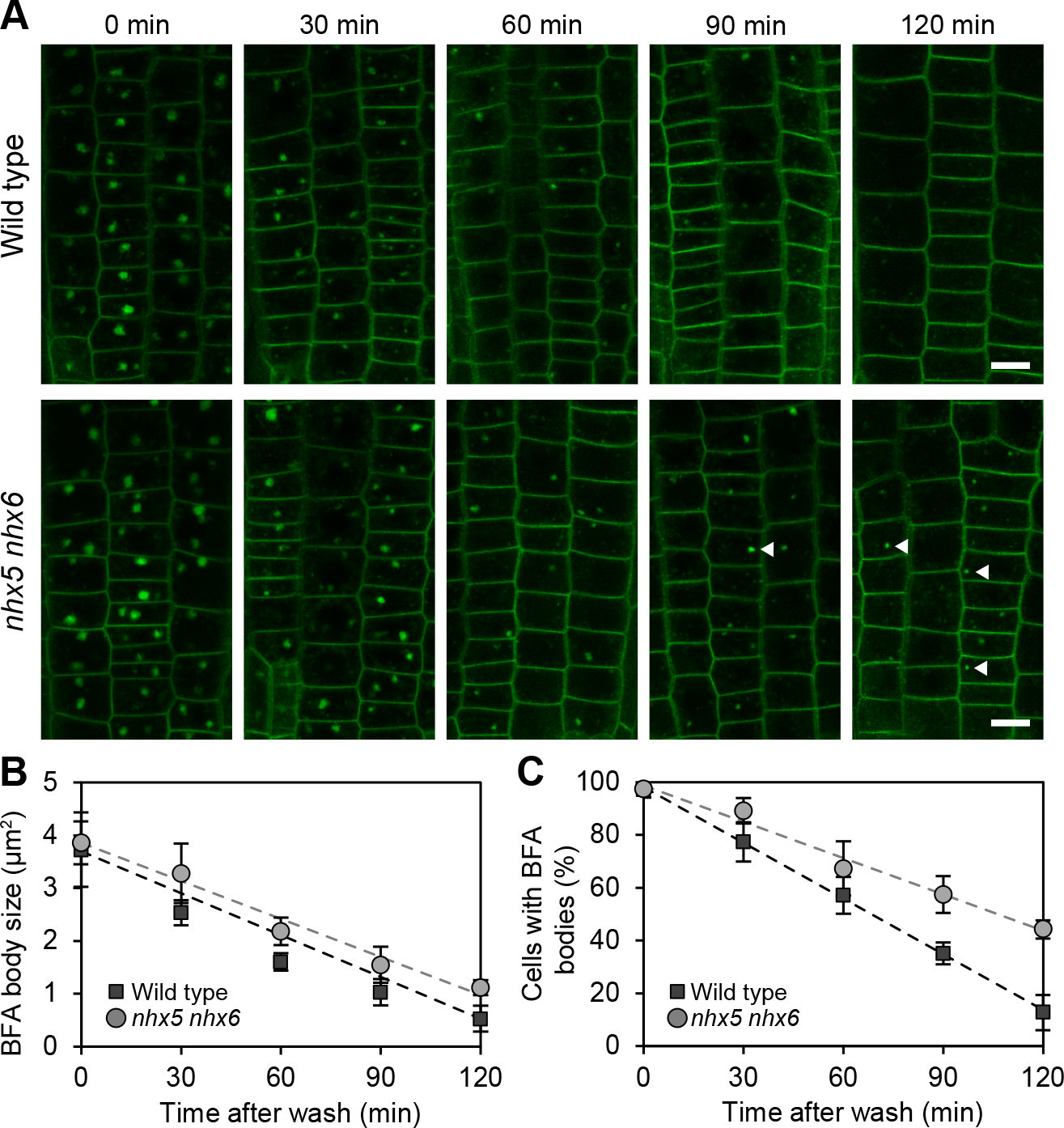
BRI1 recycling from the TGN/EE is reduced in *nhx5 nhx6*. (A) Confocal microscopy analysis of BRI1-GFP recycling after Brefeldin A (BFA) washout. Roots from 7 day old Arabidopsis seedlings were pre-treated with Cycloheximide (CHX) for 1hr followed by treatment with (BFA) and CHX for 1 hr, and washed in CHX for the indicated times. BFA bodies are still present in *nhx5 nhx6* cells after 90 and 120 mins washout (arrowheads). Scale bar = 10 μm. (B and C) Quantification of BRI1-GFP BFA body size after washout (B), and proportion of cells containing BFA bodies after washout (C). ≥ 15 cells from ≥ 3 individual plants were analysed for each time point. Data is means ± S.D.

As dysregulation of TGN/EE V-ATPase activity has been implicated in defects in protein secretion and delivery of proteins to the plasma membrane (Dettmer et al., 2006; Luo et al., 2015), we questioned whether these pathways are sensitive to general pH disruptions and could be similarly disturbed in *nhx5 nhx6* mutants. We first quantified steady state levels of BRI1-GFP at the plasma membrane in wild type and *nhx5 nhx6* roots but found no significant differences in fluorescence levels. Similarly, fluorescence recovery after photobleaching (FRAP) experiments revealed no clear difference in recovery of BRI1-GFP to the plasma membrane in *nhx5 nhx6* (Fig. 3C-D), suggesting that the delivery of newly synthesised BRI1-GFP to the plasma membrane is not impaired in *nhx5 nhx6*. We also assessed endocytic uptake of the endocytic tracer dye FM5-95 (a FM4-64 analogue) to examine whether endocytosis may be disrupted in *nhx5 nhx6* (Fig. S3). Similarly, we found no clear defect in endocytic uptake of FM4-64 dye in *nhx5 nhx6* cells. Taken together, these findings indicate that NHX5 and NHX6 activity is important for TGN/EE function and recycling of BRI1-GFP, but not for its synthesis and delivery to the plasma membrane.

**Figure 3.**
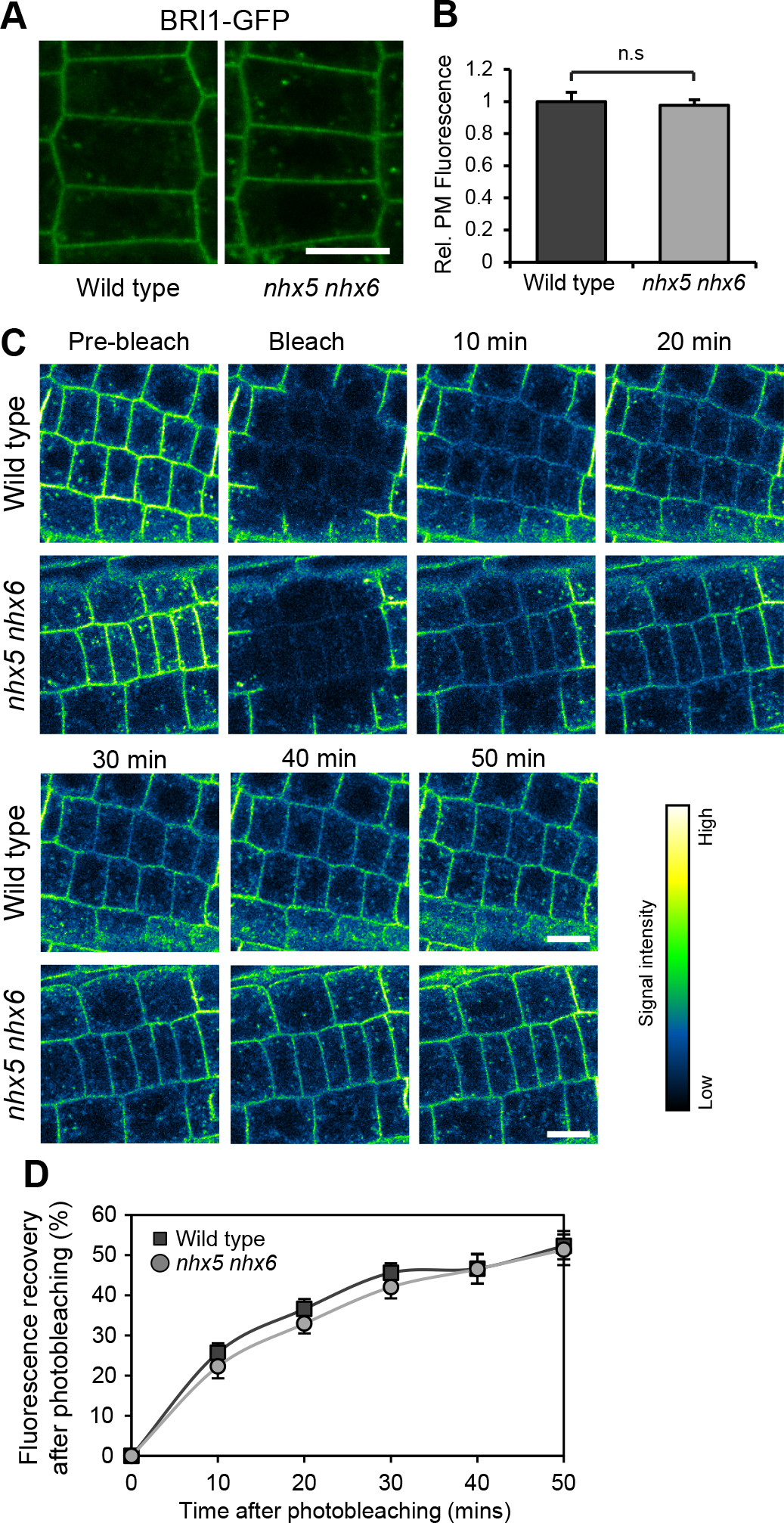
BRI1 secretion to the plasma membrane is unaffected in *nhx5 nhx6*. (A) Confocal microscopy images of root epidermal cells expressing BRI1-GFP in wild type and *nhx5 nhx6*. (B) Quantification of relative plasma membrane fluorescence. Data is means ± s.e.m., n.s.; not significant. ≥ 5 cells from ≥ 6 roots were analysed for each genotype. (C) Fluorescence recovery after photobleaching (FRAP) in seedlings expressing BRI1-GFP. Root epidermal cells were imaged pre-bleach, after bleaching with the 488nm argon laser, and at time intervals as indicated. (D) Quantification of BRI1-GFP plasma membrane fluorescence after bleaching. Data is means ± SE, n ≥ 4 cells per time point. No significant differences in rate of fluorescence recovery were observed between wild type and *nhx5 nhx6*. Scale bars = 10 μm.

### NHX5 and NHX6 antiporter activity is required for wortmannin induced swelling of MVBs

As NHX5 and NHX6 have been reported to localise to the late endosome/multi-vesicular body (LE/MVB) (Reguera et al., 2015), we questioned whether trafficking or function at the LE/MVB could be affected in *nhx5 nhx6* mutants. Wortmannin inhibits phosphatidylinositol 3-kinases (PI3K) and PI4K at high concentrations (>30 µM) and has been used extensively to investigate late endosome trafficking in plants (Jaillais et al., 2006; Vermeer et al., 2006). We examined root cells expressing the Rab5 GTPase YFP-ARA7 (RABF2b) as a LE/MVB marker. In wild type cells wortmannin induced enlarged ring-like structures (Fig. 4A) which were associated, but not completely fused with TGN/EE labelled with the SNARE VTI12 (Fig S4). Surprisingly, while similar number of wortmannin bodies were present in *nhx5 nhx6* cells compared to wild type, they were significantly smaller and denser, and did not exhibit a characteristic ring-like shape (Fig. 4A-B). To test if these wortmannin induced fusion defects were dependent on the antiporter activity of NHX6, we generated an antiporter inactive NHX6 by mutating the highly conserved acidic Asp194 residue which is critical for Na^+^ and H^+^ ion binding to Asn (D194N) (Lee et al., 2013; Wang et al., 2015), and fused it to EGFP (Fig. S5A). This mutated NHX6_D194N_-EGFP reporter did not rescue growth impairment in *nhx5 nhx6* (Fig. S5B), but localised correctly to core and peripheral BFA compartments as previously reported with functional NHX5/6 (Fig. S5C) (Bassil et al., 2011a). Wortmannin treated cells expressing NHX6_D194N_-EGFP had smaller wortmannin bodies in the *nhx5 nhx6* background compared to control seedlings (Fig. 4A-B), suggesting that NHX antiporter activity was required for swelling of LE/MVBs in response to wortmannin.

**Figure 4.**
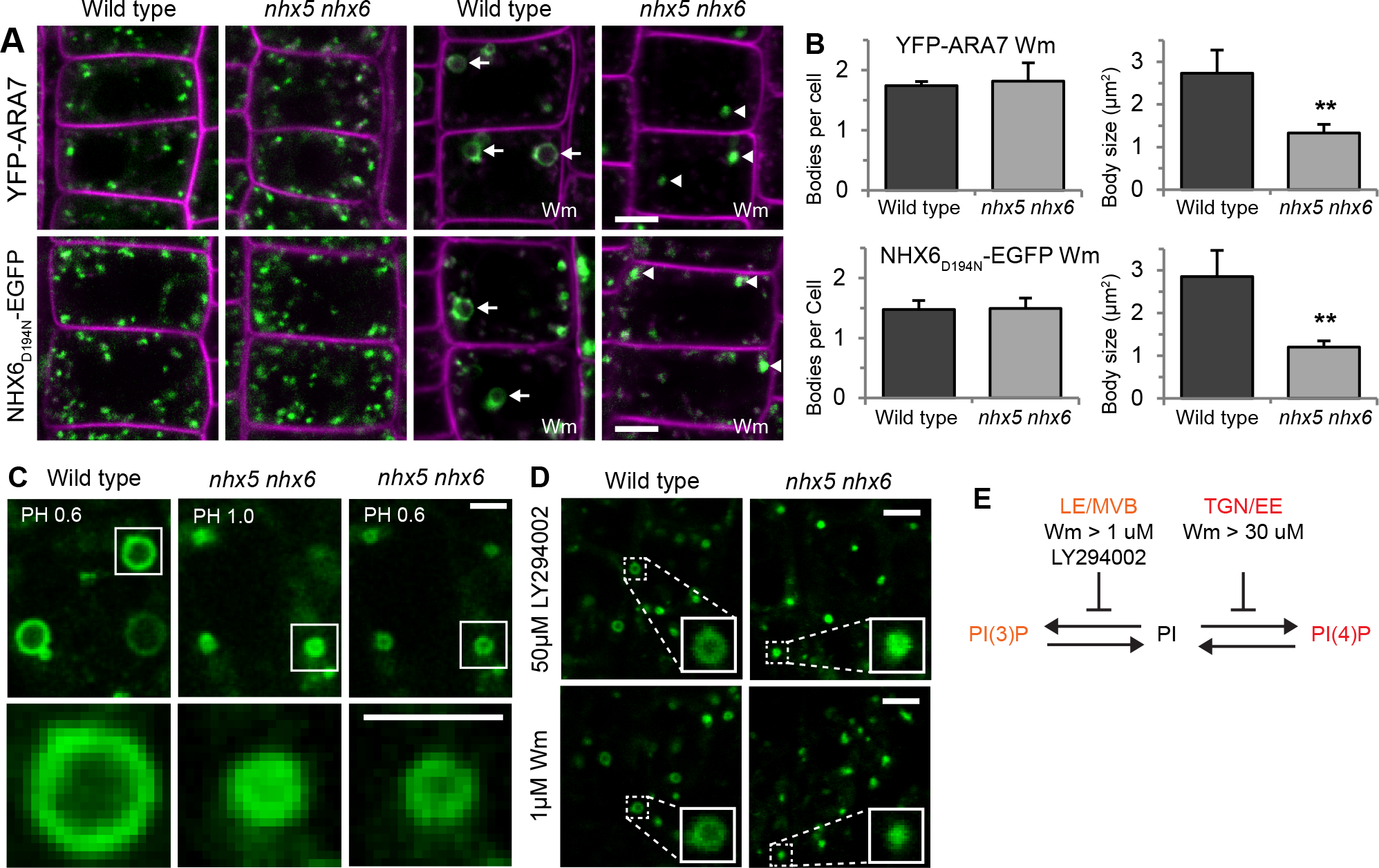
Endosomal NHX antiporter activity is required for Wortmannin induced LE/MVB swelling. (A) Confocal microscopy analysis of root epidermal cells from plants expressing YFP-ARA7 (LE/MVB marker), and the antiporter inactive NHX6_D194N_-EGFP. Cells were mock treated (0.1% DMSO) or treated with 33 μM wortmannin (Wm) for 90 minutes. Wortmannin inhibits late endosome trafficking and induced homotypic fusion of MVBs into large bodies with a ring morphology in wild type (arrows), compared to smaller, dense bodies in *nhx5 nhx6* (arrowheads). Plasma membrane is counterstained with FM5-95 in magenta. (B) Quantification of wortmannin bodies from (A). Data is means ± SD, ** p <0.01; t-test. ≥ 9 cells from ≥ 4 independent roots were analysed for each genotype. (C) Wortmannin induced bodies in *nhx5 nhx6* are resolvable through confocal imaging with a reduced pinhole. Root cap cells treated with 33 μM wortmannin and imaged with a 0.6 Airy Unit pinhole (PH) show a hollow ring structure. (D) Response of root cap cells expressing YFP-ARA7 to 50 μM LY294002 or 1 μM wortmannin treatment. Inset shows an enlarged MVB/LE body. (E) Schematic of inhibitors used to interrupt phosphoinositide conversion. LY294002 and 1 μM wortmannin specifically inhibit PI3K conversion of PI to PI(3)P, while concentrations above 30 μM Wm also inhibit PI4K conversion of PI to PI(4)P. Scale bars = 5 μm (A, C), 2 μm (D).

Next, we questioned whether the structure of the small wortmannin bodies in *nhx5 nhx6* cells were different from bodies in wild type cells. We imaged wortmannin treated cells expressing YFP-ARA7 with a reduced confocal pinhole diameter to increase spatial resolution in x, y and z dimensions. Under reduced pinhole settings wortmannin bodies in *nhx5 nhx6* root cells were resolvable and had a clear ring-like structure similar to wild type cells (Fig. 4C), suggesting they shared a similar structure. Since wortmannin inhibits both PI3K and PI4K at high concentrations, we assessed whether *nhx5 nhx6* swelling insensitivity was present during PI3K specific inhibition. Specifically targeting PI3K pathways by using the inhibitor LY294002 or with low concentrations of wortmannin (Fujimoto et al., 2015; Simon et al., 2016; Takáč et al., 2013), also caused similar fusion of MVBs into smaller, more densely labelled bodies in *nhx5 nhx6* cells (Fig. 4D). This finding indicates that wortmannin and LY294002 induced swelling is caused by inhibition of PI(3)P on the LE/MVB (Fig. 4E). Taken together, this data shows that along with inducing LE/MVB fusion, wortmannin induces rapid osmotic swelling of fused MVB bodies, consistent with data showing wortmannin causes alkalinisation of LE/MVBs (Martinière et al., 2013). Thus, the insensitivity to swelling in *nhx5 nhx6* are a consequence of the lack of normal ion or pH regulation mediated by NHX5 and NHX6 antiporter activity.

### NHX5 and NHX6 do not regulate root vacuole morphology

In plants, the primary vacuolar transport pathway is marked by the co-ordinated sequential activity of the Rab GTPases Rab5 (RabF) and Rab7 (RabG) (Cui et al., 2014; Cui et al., 2016). This pathway is also implicated in the transport of storage proteins to the protein storage vacuole (PSV) in seeds (Ebine et al., 2014; Singh et al., 2014), which has shown to be disrupted in *nhx5 nhx6* (Ashnest et al., 2015; Reguera et al., 2015). The constitutively active GTP-bound mutant of Rab5 (ARA7QL) transits through the LE/MVB to the tonoplast (Ebine et al., 2011), allowing us to assess whether Rab5 recruitment or maturation could be affected in *nhx5 nhx6* root cells. We observed similar localisation of ARA7QL to the LE/MVB and tonoplast in wild type and *nhx5 nhx6* cells, however *nhx5 nhx6* cells were insensitivity to swelling induced by wortmannin (Fig. S6A), similar to our findings using the unmutated YFP-ARA7 reporter (Fig. 4A). As endosomal NHX type antiporter activity has been previously reported to be implicated in vacuolar trafficking and fusion in the yeast *nhx1* mutant (Qiu and Fratti, 2010), we assessed vacuole morphology in *nhx5 nhx6* root cells, but found no clear abnormalities in vacuolar structure or response to wortmannin (Fig. S5B-C). Together these results suggest NHX5 and NHX6 are not important for vacuole function in root tissue.

### *nhx5 nhx6* endosomes have reduced sensitivity to monensin

Since NHXs regulate Na^+^ and K^+^ accumulation in the endosomal lumen, we questioned whether the reduced swelling of LE/MVB induced by wortmannin in *nhx5 nhx6* could be due to an altered ionic composition in the LE/MVB lumen. The monovalent cation ionophore monensin induces rapid osmotic swelling of *trans*-Golgi stacks and TGN/EE through transport of Na^+^/K^+^ for H^+^ across the endomembrane (Zhang et al., 1993). Consistent with this data, we observed rapid swelling TGN/EE, but not of LE/MVB in root cells (Fig. 5A). Next, we assessed the pH of TGN/EE after monensin treatment using the TGN/EE localised ratiometric pH sensor SYP61-pHusion (Luo et al., 2015). TGN/EE of monensin treated root cells were less acidic than untreated cells (Fig. 5B), consistent with monensin induced luminal import of Na^+^/K^+^ in exchange for H^+^. Furthermore, we assessed the pH of TGN/EE of *nhx5 nhx6* root cells and found a ~0.5 pH reduction (acidification) of TGN/EE (Fig. 5C), similar to previous reports in Arabidopsis protoplasts (Reguera et al., 2015).

**Figure 5.**
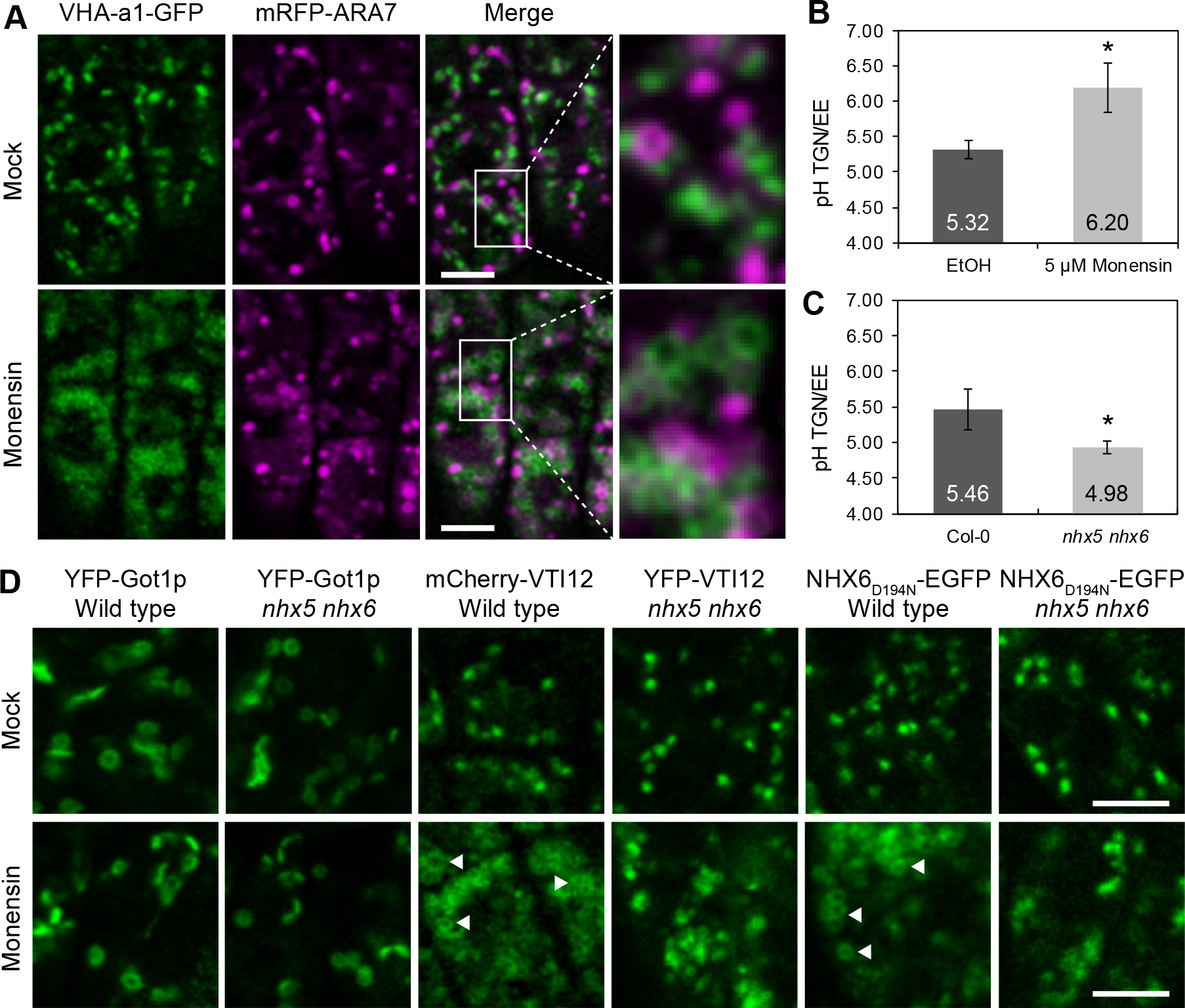
*nhx5 nhx6* TGN/EE are partially insensitive to monensin induced swelling. (A) Confocal microscopy images of root cap cells treated with 10 μM monensin for 30 minutes. The ionophore monensin induces swelling of TGN/EE (VHA-a1-GFP), but not of LE/MVBs (mRFP-ARA7); inset shows magnified view of enlarged TGN/EE. (B) pH of TGN/EE from seedlings expressing the ratiometric pH sensor SYP61-pHusion after treatment with 5 μM monensin for 30 minutes. (C) pH of TGN/EE from wild type and *nhx5-1 nhx6-1* seedlings expressing SYP61-pHusion. Data from B-C represent means ± SD of three individual experiments from n = 15 seedlings. * p < 0.05; Student’s t-test. (D) Arabidopsis root cap cells following 30 minutes 10 μM monensin treatment, or mock control (0.1% EtOH). Swelling of Golgi (YFP-Got1p) was not detected. TGN/EE (YFP-VTI12) showed aggregation and enlargement of individual endosomes (arrowheads) which was less apparent in *nhx5 nhx6*. Similar results were obtained with the NHX6 antiporter inactive marker NHX6_D194N_-EGFP. Scale bars, 5 μm.

Given the increased acidity of *nhx5 nhx6* TGN/EE, we questioned whether *nhx5 nhx6* root cells could be hypersensitive to swelling induced by monensin, given they produce a stronger proton gradient for monensin to act upon. We therefore assessed the response to monensin using our established Golgi and TGN/EE markers in *nhx5 nhx6*. We could not detect any significant swelling of Golgi in either wild type or *nhx5 nhx6* monensin treated root cells (Fig. 5D), likely as the *trans* most Golgi stack only swells slightly upon monensin treatment (Zhang et al., 1993). TGN/EE in monensin treated roots showed clear clustering and swelling in wild type, however in *nhx5 nhx6* only minor swelling was present despite clustering of TGN/EE, suggesting these TGN/EE had reduced capacity to swell (Fig. 5D). Furthermore, similar results were obtained using NHX6_D194N_-EGFP as a marker for NHX6 activity in wild type and *nhx5 nhx6* backgrounds (Fig. 5D). Together, these results suggest that in *nhx5 nhx6* cells TGN/EE have reduced or slowed osmotic swelling induced by monensin, likely originating from a disruption to intraluminal ion (K^+^) balance in these endosomes.

### The C-terminal cytosolic tail of NHX6 mediates its localisation to the MVB

We previously reported that the cytosolic tail of NHX6 interacts with the retromer component SNX1 (Ashnest et al., 2015), however it is unclear what functional significance this interaction has, and whether the cytosolic tail mediates other functions. To investigate this, we generated a partial tail truncation of NHX6 lacking a short 28 amino acid region of the C-terminus fused to YFP (Fig. 6A), as a previous truncation lacking the entire C-terminal tail did not show stable expression (Ashnest et al., 2015). This NHX6_1-507_-YFP construct completely complemented the dwarf phenotype of *nhx5 nhx6* plants (Fig. 6B), suggesting that this distal region of tail was not essential for NHX6 antiporter activity.

**Figure 6.**
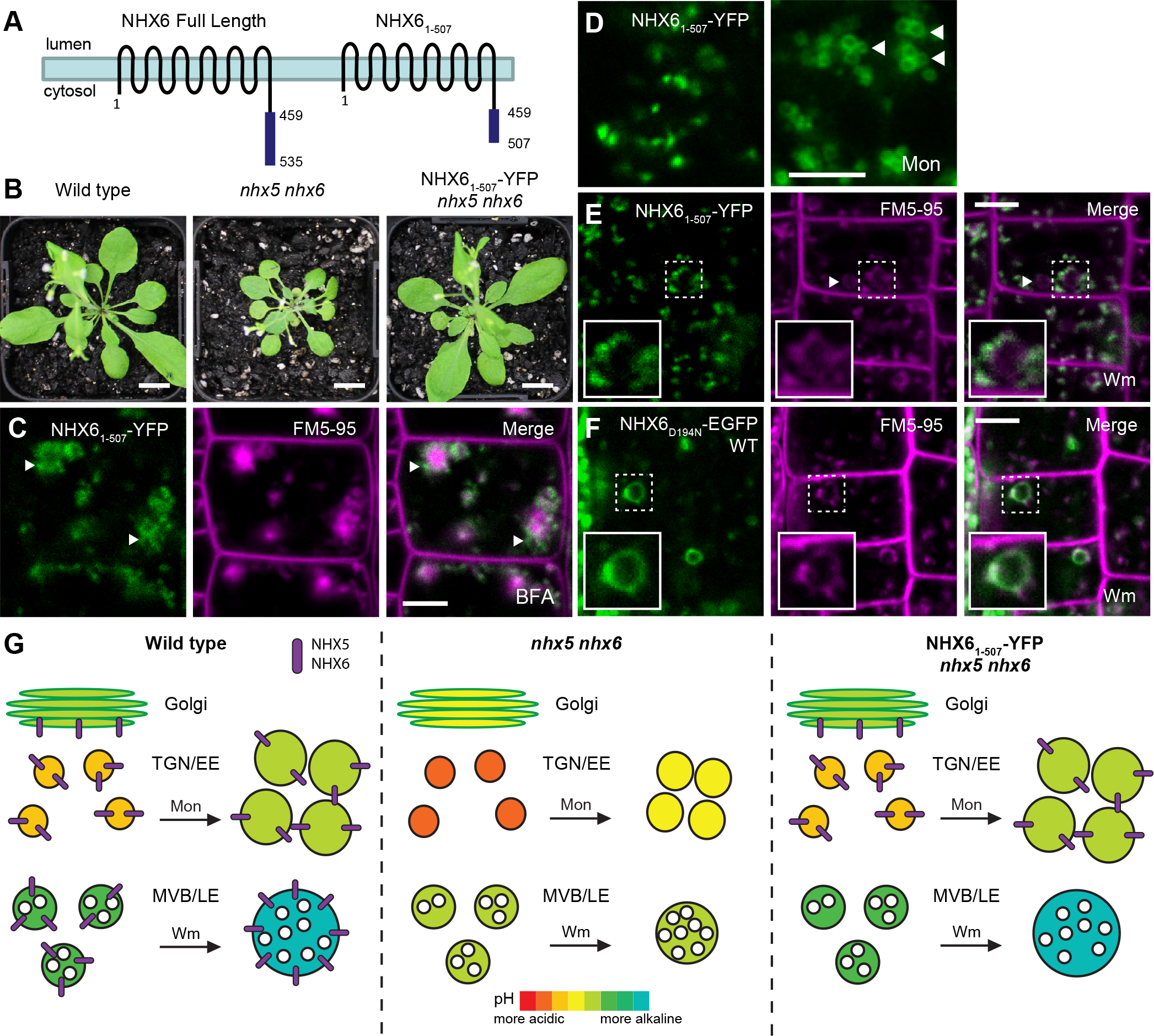
The distal tail of NHX6 is required for localisation to the MVB. (A) Structure of the NHX6_1-507_ deletion mutant construct. (B) Complementation of the *nhx5 nhx6* dwarf phenotype by NHX6_1-507_-YFP in 28 day old Arabidopsis plants. (C) Confocal images of root epidermal cells in *nhx5-2 nhx6-3*/NHX6_1-507_-YFP treated with BFA for 60 minutes. Note the fluorescence signal in core and peripheral (arrowheads) BFA body regions, corresponding to TGN/EE and Golgi respectively. (D) Root cap cells from NHX6_1-507_-YFP seedlings following mock treatment (0.1% DMSO) or treatment with 10 μM monensin (Mon) for 30 minutes. Note the enlargement of TGN/EE (arrowheads). (E-F) Root epidermal cells from NHX6_1-507_-YFP or WT/NHX6_D194N_-EGFP seedlings treated with to 33 μM wortmannin (Wm). Arrowheads indicate lack of NHX6_1-507_-YFP labelling in Wm bodies. Partial labelling of NHX6_1-507_-YFP to a Wm body (inset), compared to complete labelling in NHX6_D194N_-EGFP. FM5-95 counterstain (magenta) labels the plasma membrane and endocytic compartments. (G) Schematic of monensin and wortmannin swelling phenotypes. In wild type cells the Golgi, TGN/EE, and MVB/LE in swell upon monensin and wortmannin/LY294002 treatment due to Na^+^/K^+^ transport for H^+^, resulting in alkalinisation. In *nhx5 nhx6* endosomal compartments are more acidic, and we propose have lower luminal Na^+^/K^+^, leading to only minor swelling with monensin, and fusion but no swelling with wortmannin. NHX6 activity at the Golgi and TGN/EE, but not MVB in NHX6_1-507_-YFP/*nhx5 nhx6* is sufficient to restore both monensin and wortmannin insensitivity phenotypes.

We inferred the localisation of this tail truncated, but antiporter active protein through pharmacological inhibitor experiments. NHX6_1-507_-YFP was sensitive to BFA, localising to core and peripheral BFA compartments, corresponding to the TGN/EE and Golgi respectively (Fig 6C) (Naramoto et al., 2014; Richter et al., 2007), similar to full length NHX6 (Fig. S5). Moreover, NHX6_1-507_-YFP was able to restore monensin insensitivity of *nhx5 nhx6*, confirming antiporter function at the TGN/EE. However, in response to inhibition of late endosomal trafficking by wortmannin, NHX6_1-507_-YFP was only partially associated with wortmannin bodies unlike the full length NHX6 (Fig. 6E-F), a response typical of slow fusion of TGN/EE markers with the wortmannin body (Fig. S4), and suggested NHX6_1-507_-YFP may be absent from the MVB. Surprisingly, in these wortmannin treated seedlings, the enlarged wortmannin bodies were morphologically similar to wild type, indicating NHX6_1-507_-YFP can also restore the wortmannin induced swelling insensitivity in *nhx5 nhx6* (Fig. 6E). As the LE/MVB matures from TGN/EE derived compartments (Scheuring et al., 2011), NHX6_1-507_-YFP antiporter activity at the Golgi and TGN/EE may be sufficient to maintain relatively normal pH in LE/MVB compartments, and thus may restore normal MVB function (Fig. 6G). Taken together, this data indicates that the distal region of the cytosolic tail is important for NHX6 localisation to the LE/MVB, but NHX6 activity at the MVB is not essential for wortmannin induced MVB swelling.

## Discussion

In this study we investigated the function of NHX5 and NHX6 in endomembrane trafficking and ion homeostasis in *Arabidopsis thaliana*. Previous studies identified that NHX5 and NHX6 localise to the Golgi, TGN/EE, and LE/MVB, and play roles in soluble cargo trafficking in seeds (Bassil et al., 2011a; Reguera et al., 2015). Here we show that NHX5 and NHX6 play additional roles in important endomembrane processes including endosomal motility and trafficking at the TGN/EE. Furthermore, we report that monensin and wortmannin induces osmotic swelling of endosomes which is affected by endosomal ion homeostasis maintained by NHX5 and NHX6. Our results demonstrate that NHX antiporters are important regulators of endomembrane pH and ion homeostasis required for efficient endomembrane trafficking.

### The NHX6 C-terminal cytosolic tail is essential for localisation to the MVB

We previously reported that the cytosolic tail of NHX6 was essential for NHX6 activity, protein stability, and mediates an interaction with the retromer component SORTING NEXIN 1 (SNX1) (Ashnest et al., 2015). Here, we generated a partial tail deletion construct (NHX6_1-507_-YFP) which appeared to be fully functional as it could completely restore the dwarf phenotype of *nhx5 nhx6* mutants. Inference of NHX6_1-507_-YFP localisation by pharmacological inhibition revealed that this distal region (amino acids 508-535) of the cytosolic tail is required for mediating NHX6 localisation to the LE/MVB. Interestingly, the cytosolic tail of the K^+^ transporter CHX17 has also been reported to mediate its localisation to the LE/MVB (Chanroj et al., 2013), however lack of significant sequence similarity between CHX17 and NHX5/6 suggests there is unlikely to be a strict sequence based sorting mechanism mediating endosomal cation/proton exchanger localisation. It is unclear whether a conserved amino acid motif in this region, and/or other signals in the transmembrane domain may mediate the sorting of NHX5 and NHX6 or other endosomal transporters to the LE/MVB.

### Regulation of endosomal ion composition by NHX5 and NHX6

Here we identified a novel mechanism of swelling of late endosomes induced by wortmannin which shares a striking resemblance to osmotic swelling of TGN/EE induced by the cation ionophore monensin. While wortmannin is well described to cause enlargement of LE/MVBs from homotypic membrane fusion of MVBs (Wang et al., 2009; Zheng et al., 2014), the typical size of these wortmannin bodies is much larger than can be achieved through membrane fusion alone. Monensin induced TGN/EE swelling occurs through Na^+^/K^+^ transport into the endosomal lumen across a proton gradient (Zhang et al., 1993), resulting in luminal alkalinisation of endosomes consistent with our reported pH measurements. Similarly, wortmannin treatment induces alkalinisation of the LE/MVB (Martinière et al., 2013), and suggests swelling induced by wortmannin may be a consequence of rapid luminal cation (eg: K^+^) import.

It is currently not known how the presumed cation import is induced by wortmannin in fused late endosomes. Since NHX6_1-507_-YFP activity at the TGN/EE but not LE/MVB could restore wortmannin swelling insensitivity, this suggests that potential K^+^ influx occurs independently of endosomal NHX5 and NHX6 activity. This might involve ion import into the MVB lumen by the wortmannin sensitive K^+^/H^+^ transporter AtCHX17 (Chanroj et al., 2013), or could be a general consequence from rapid homotypic fusion of LE/MVB. We speculate that inhibition of PI3K through wortmannin or LY294002 and the resulting loss of LE/MVB membrane identity could lead to de-repression of cation transporters, and therefore facilitate uncontrolled ion influx and induce osmotic swelling. Investigation of changes in endosomal K^+^ concentrations *in vivo* using recently established genetically encoded fluorescent K^+^ probes may facilitate greater understanding of cation compositions and dynamics of endosomal compartments (Bischof et al., 2017).

Our results demonstrate that *nhx5 nhx6* seedlings have reduced swelling of TGN/EE compartments upon monensin treatment. As plant NHXs function in intracellular internalisation of Na^+^ and K^+^ (Bassil et al., 2011a; Bassil et al., 2011b), endosomes in *nhx5 nhx6* likely have reduced concentrations of luminal K^+^ ions. Thus, while monensin retains functional activity as a cation importer and causes aggregation of TGN/EE in *nhx5 nhx6*, osmotic swelling occurs more slowly due to the reduced luminal cation concentration at the TGN/EE (Fig. 6G). Furthermore, the insensitivity of *nhx5 nhx6* late endosomes to swelling by wortmannin treatment appears to occur due to a similar cation balance at the late endosome. As the LE/MVB matures from the TGN/EE by budding (Scheuring et al., 2011), these compartments will likely share a similar ion composition as its source membrane (TGN/EE) which would explain why NHX6_1-507_-YFP could restore wortmannin insensitivity despite not localising to the LE/MVB itself. Accordingly, NHX activity at the MVB appears largely inconsequential for maintaining ion balance at the MVB.

### NHX5 and NHX6 are important for endosome motility

Here we identified that Golgi and TGN/EE motility is significantly reduced in *nhx5 nhx6* hypocotyl cells, and that the behaviour of TGN/EE is altered, suggestive of a reduction in the ability of TGN/EE to associate with the cytoskeleton. These findings are similar to reported defects of Golgi and TGN/EE motility in *det3* mutants which also have altered TGN/EE pH (Luo et al., 2015), however the functional consequences of a reduction in endosome motility is not clear. Previous reports indicate that root hair cell growth is significantly reduced in *nhx5 nhx6* knockouts (Bassil et al., 2011a). As tip directed growth of root hair cells relies on rapid endosomal transport along actin filaments (Szymanski and Staiger, 2018), we speculate that the reduced endosomal motility in *nhx5 nhx6* may lead to slowed endosomal transport to the growing cell tip and consequently inhibit the rate of cell expansion. Similarly, reduced endosome motility in the hypocotyl could explain the reduction in hypocotyl cell elongation in both *nhx5 nhx6* and *det3* mutants, and suggests a pH sensitive mechanism governs endosome-cytoskeleton association.

### NHX5 and NHX6 play roles in recycling at the TGN/EE

Pharmacological BFA and FRAP experiments indicate that *nhx5 nhx6* have defects in BRI1-GFP recycling from the TGN/EE, but not in general BRI1 secretion to the plasma membrane. These results contrast with findings in *det3* V-ATPase mutants which have both impaired secretion and recycling of BRI1 (Luo et al., 2015). Given that *det3* mutants have alkalinised TGN/EE due to reduced V-ATPase activity, while *nhx5 nhx6* have hyper-acidified TGN/EE, these findings suggest that maintaining correct homeostasis of TGN/EE pH is essential for functional protein recycling from the TGN/EE. This idea is consistent with *nhx5 nhx6* mutants showing similar hypersensitivity to Concanamycin A treatment as *det3*, implying a defect at the TGN/EE. Interestingly, we previously reported that the transport of polar localised auxin carrier proteins PIN1-GFP and PIN2-GFP from the TGN/EE was not significantly impaired in *nhx5 nhx6* root cells (Dragwidge et al., 2018). This finding is consistent with emerging evidence that the TGN/EE is composed of functionally segregated domains that sort distinct membrane cargoes for delivery to the plasma membrane (Li et al., 2016; Singh and Jürgens, 2017). Thus, correct regulation of TGN/EE pH homeostasis may be more critical for certain TGN/EE trafficking pathways than others.

How NHX5 and NHX6 affect trafficking at the TGN/EE is unknown. In animals and *Drosophila*, NHX and V-ATPase activity has been shown to effect electrostatics at the cytosolic surface of endomembranes which can affect membrane signalling or protein recruitment. Specifically, endosomal V-ATPase activity is required for recruitment of the small GTPase Arf6 In mammalian cells (Hurtado-Lorenzo et al., 2006), while pH- and charge-dependent Wnt signalling at the plasma membrane is regulated by Nhe2 activity in Drosophila (Simons et al., 2009). Thus, potential disruptions to endomembrane electrostatics in *nhx5 nhx6* may influence the charge-dependent recruitment of small GTPases or Arfs involved in protein trafficking or recycling. Further investigation of endosomal trafficking pathways through selective trafficking inhibitors such as Endosidin compounds may uncover a clearer understanding of the specific trafficking pathways which require NHX5 and NHX6 activity (Hicks and Raikhel, 2010; Li et al., 2016).

In conclusion, we have shown that endosomal pH and ion regulation by plant NHXs is important for functional endosomal behaviour. We demonstrate that NHX5 and NHX6 activity is necessary for functional Golgi and TGN/EE motility, protein recycling at the TGN/EE, and for regulation of endomembrane ion balance. This work sheds light on the complex nature of the plant endomembrane system and demonstrates the importance of regulation of endomembrane ion and pH homeostasis.

## Materials and Methods

### Plant material and growth conditions

*Arabidopsis thaliana* lines were all in the Columbia-0 (Col-0) accession background. Plant lines used have been previously described, *nhx5-1 nhx6-1* (Bassil et al., 2011a), *nhx5-2 nhx6-3* (Ashnest et al., 2015), *det3* (Schumacher et al., 1999), pBRI1::BRI1–GFP (Geldner et al., 2007), GFP-ARA7Q69L (Ebine et al., 2011), pUBQ10::xFP-RabF2b (Wave2Y/R), pUBQ10::YFP-Got1p (Wave18Y), pUBQ10::xFP-VAMP711 (Wave 9Y/R), pUBQ10::YFP-

VTI12 (Wave13Y) (Geldner et al., 2009), VHA-a1-GFP × mRFP-ARA7 and 2xFYVE-GFP x VHA-a1-mRFP (Singh et al., 2014). The *nhx5-2 nhx6-3* allele combination was used for all experiments except for the SYP61-pHusion pH measurements and ConcA hypocotyl assay where *nhx5-1 nhx6-1* was used. BRI1-GFP in *nhx5 nhx6* was generated by crossing BRI1-GFP into NHX5 / *nhx5 nhx6* plants, and *nhx5 nhx6* plants homozygous for BRI-GFP were identified in the following generations. The *nhx5 nhx6* lines expressing other fluorescent markers were obtained by floral dripping (Martinez-Trujillo et al., 2004) using Agrobacterium *tumefaciens* GV3101 strain cultures containing the given constructs.

Seeds were surface sterilized with 70% ethanol for 5 minutes, 10% bleach for 5 minutes, and washed three times in ddH_2_O and grown on ½ strength Murashige and Skoog (½ MS) medium containing 1.0% (w/v) agar, pH 5.8 without sucrose unless indicated. Seedlings were stratified for 48 hours at 4°C in the dark and grown in a 16hr light/8hr dark photoperiod at 22°C at 100 umol m^−2^s^−1^ under cool-white fluorescent lights.

For live cell microscopy and chemical treatments, seedlings were grown vertically on ½ MS plates with 1.0% (w/v) agar without sucrose. For time lapse motility assays, etiolated seedlings were grown at 22 °C in the dark for 4 days.

### Construct Generation

For all cloning procedures, the Gateway recombination system (Thermo-Fisher) was used. NHX6_1-507_-YFP was generated by amplifying amino acids 1-507 of NHX6 from a cloned NHX6 ORF plasmid template using NHX6FS and NHX6 507R primers with iProof polymerase. This fragment was cloned into pENTR-D-TOPO, sequenced, and recombined into pEarlyGate 101 (Earley et al., 2006) to generate 35S:NHX6_1-507_-YFP. NHX6_D194N_-EGFP was generated through site directed mutagenesis using partially overlapping primers. D194N-F1 and D194N-R1 primers were used to amplify from a NHX6-EGFP pENTR-D-TOPO template using PfuUltra high-fidelity DNA polymerase, then LR recombined into pEarlyGate 100 to generate 35S:NHX6_D194N_-EGFP.

### Confocal microscopy and drug treatments

Images were obtained on a Leica SP5 or Zeiss LSM 780 laser scanning confocal microscope using a C-Apochromat 40x/1.3 water objective with 2x digital zoom at 1024×1024x pixels per image. Excitation and emission detection settings were as follows: mCerulean 458 nm/460-520 nm; GFP/YFP 488 nm/490-560 nm; mRFP/mCherry 561 nm/565-630 nm; FM5-95 561 nm/565-650 nm. For all quantification experiments, identical settings were used to acquire each image. Chemical stock solutions were made in DMSO at the following concentrations - BFA 50 mM (Sigma-Aldrich), CHX 50 mM (Sigma-Aldrich), FM5-95 4 mM (FM4-64 analogue, ThermoFisher), Wortmannin 33 mM (LC Laboratories), LY294002 50 mM (Sigma-Aldrich), 1 mM Concanamycin A (Santa Cruz Biotechnology). Monensin stock was made in Ethanol at 10mM (Santa Cruz Biotechnology). An equal concentration of DMSO or ethanol was used in control treatments. For drug treatments, 6-7 day old seedlings were incubated in 6 well plates for 60 minutes for BFA treatments, 90 minutes for wortmannin and LY294002 treatments, or 30 minutes for monensin treatments.

For quantification of BRI1-GFP BFA bodies, images were of 3-4 slice Z-stacks with 2 μm spacing, from which maximum intensity projections were generated. BFA body size was quantified using Fiji (Schindelin et al., 2012) based on ImageJ v1.48g, with automated thresholding and the Analyse Particles tool, using a minimum size of 0.7 μm^2^, and circularity of ≥ 0.3. At least 15 epidermal cells were counted from each root. For quantification of wortmannin bodies, circular regions of interest (ROIs) were manually drawn and body area was measured using ImageJ. For analysis of bodies in NHX6_1-507_-YFP and YFP-VTI-12 lines only clearly fused bodies ≥ 1 μm^2^ were used for analysis. For quantification of BRI1-GFP plasma membrane fluorescence, plasma membrane regions from ≥ 6 cells per root were quantified using fixed ROIs, and mean fluorescence was calculated after subtracting background fluorescence. For FM internalisation, 7 day old seedlings were incubated in 2 μM FM5-95 in 6 well plates on ice for 5 minutes, washed in ½ MS twice, and incubated at room temperature for 5 minutes before confocal imaging. For ConcA hypocotyl length measurements, etiolated seedlings were scanned and measured using the “segmented line” tool in ImageJ. For monensin images and low pinhole images, a gaussian blur filter was applied with 0.6 sigma.

### Fluorescence recovery after photobleaching (FRAP)

FRAP analysis was performed on a Zeiss LSM 780 laser confocal microscope using a 40x/1.2 water objective. 7 day old seedlings were transferred from plates onto single well Lab-Tek™ Chambered Coverglass slides, and a thin agar slice was placed on top.

*Arabidopsis* root epidermal cells were bleached using a 488 nm argon laser at 100% power for 60 seconds with a circular 30 μm diameter ROI. Images were acquired using 512 × 512 pixel resolution pre-bleach at 0 min, 10 min, 20 min, 30 min, 40 min, and 50 min after bleaching.

Image series were aligned using Stackreg and Linear Stack Alignment with SIFT in ImageJ. For analysis, a fixed ROI was used to select plasma membrane from completely bleached cells. Fluorescence values from bleached cells were normalised to fluorescence from ROIs from two unbleached cells in the same image to account for minor photobleaching during image acquisition. Plasma membrane fluorescence before bleaching was set to 100%, and directly after bleaching as 0%. Percent of fluorescence recovery after photobleaching was calculated by dividing the normalised bleached fluorescence value minus background (t = 0 min) from the pre-bleached value minus background (t = 0 min). The experiment was repeated, and similar results were obtained.

### Endosome motility imaging and analysis

Time lapse motility experiments were performed on a Zeiss Cell Observer spinning disk confocal microscope equipped with a Yokogawa CSU-X1 spinning disk and Photometrics EMCCD camera, using a 63x oil immersion objective. Four day old dark grown seedlings were placed onto single well Lab-Tek™ Chambered Coverglass (ThermoFisher) and a thin agar slice was placed on top. Epidermal cells from the upper hypocotyl were imaged at the cortical focal plane just below the plasma membrane over a 2 min period with 1 s scanning intervals. Only similarly sized cells (~900 - 1200 μm^2^) were selected for analysis to minimise variation in particle speeds due to cell size. Cytoplasmic streaming was observed to verify cell viability.

Image drift was corrected using the ImageJ plugin StackReg. Postprocessing and analysis was performed in IMARIS software v7.0 (Bitplane). Particle tracking was performed using the ‘spots’ feature with the autoregressive motion algorithm, with a max distance parameter of 10 pixels, and a gap parameter of 0. Particle tracks less than 10 seconds long were filtered and excluded from analysis. Tracks were verified manually, and misaligned tracks were realigned. Mean track speed and track straightness was calculated in IMARIS from pooled particle tracks from ≥ 5 individual seedlings. Kymographs were created in ImageJ using the MultipleKymograph plugins (http://www.embl.de/eamnet/html/body_kymograph.html).

### Vacuolar morphology

For vacuolar morphology analysis 7 day old seedlings expressing YFP-VAMP711 or RFP-VAMP711 were imaged on a Zeiss spinning disk confocal microscope using a 63x oil immersion objective. Z-stacks from Atrichoblast epidermal cells were obtained from 30-45 slices with 200 nm step size and stacked as a maximal intensity projection. Vacuolar morphology index was calculated by measuring the maximal luminal length and width in each cell in ImageJ (Löfke et al., 2015). For quantification ≥ 5 cells from ≥ 5 individual seedlings were analysed.

### TGN/EE pH measurements

pH measurements of TGN/EE using the SYP61-pHusion line was performed as previously described (Luo et al., 2015). Briefly, 6-7 day old seedlings expressing SYP61-pHusion were imaged on a Leica SP5 scanning confocal microscope with a HCX PL APO 63× 1.20 water immersion objective with GFP (ex 488, em 490-545), and mRFP (ex 561, em 600-670) settings. Ratios were calculated by dividing the average intensity of GFP/mRFP signals after background subtraction. pH was calibrated *in vivo* from buffers between pH 4.8 and 8.0 using free cytosolic pHusion, from which a sigmoidal calibration curve was obtained through the Boltzmann fit function in Origin Pro 9.1G and mapped to the corresponding measured values.

### Statistics and Software

Statistics were performed in Microsoft Excel or R v3.3.2 using R Studio. Boxplots and stripcharts were generated in R v3.3.2. Images were prepared in Illustrator (Adobe).

## Acknowledgements

We thank the ABRC, Joanne Chory, and Tomohiro Uemura for sharing published materials, and the LIMS Bioimaging Facility and Peter Lock for microscopy assistance.

## Author Contributions

J.M.D. conducted the experiments and analysed the data. S.S. performed the pH measurements and Concanamycin A experiment. J.M.D., K.S., and A.R.G. designed the experiments. J.M.D and A.R.G wrote the manuscript.

---

**Table S1.**
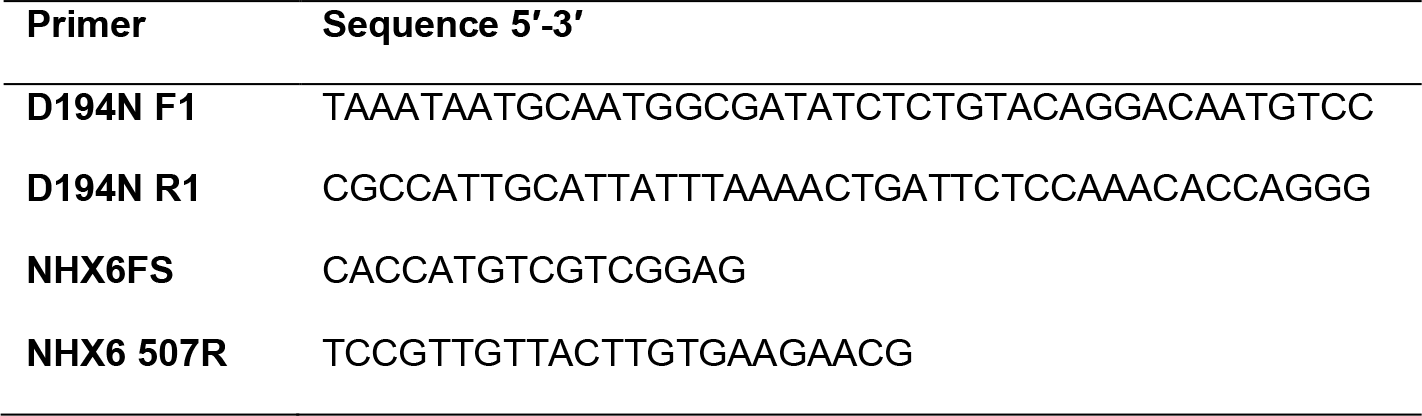
List of primers used in this study.

